# The gut microbiome is associated with cocaine behavior and predicts addiction vulnerability in adult male rats

**DOI:** 10.1101/2021.07.20.453110

**Authors:** Gregory J Suess, Jennysue Kasiah, Sierra Simpson, Molly Brennan, Dana Conlisk, Lisa Maturin, Olivier L George, Benoit Chassaing, Kyle J Frantz

**Author notes:** Correspondence should be addressed to Kyle Frantz. These authors to be designated as co-last authors (Benoit Chassaing, Kyle J Frantz). **Author contributions: GS, SS, OG, BC, and KJF** contributed to the study concept and design. **SS, MB, and LM** contributed to the behavioral data and tissue acquisition. **GS, JK, and BC** processed the microbiota samples. **GS, SS, OG, BC, and KJF** contributed to data analysis, interpretation of data, and manuscript preparation. **All authors** approve the manuscript. Conflicts of Interest: No.

## Abstract

The gut-brain axis is a bi-directional communication system through which microbial communities in the gut interact with the nervous system, perhaps influencing neuropsychiatric disorders such as drug abuse. This study used behavioral data and biological samples from the Cocaine Biobank to test the hypothesis that the gut microbiota can predict and reflect susceptibility to cocaine reinforcement. Adult male heterogenous rats were catheterized and allowed to self-administer cocaine in short-access sessions (2 hr/day, 10 days, 0.5 mg/kg per intravenous infusion), followed by progressive ratio (PR) tests, long-access sessions (6 hr/day, 14 days), and alternating blocks of PR, long-access, and footshock testing. Fecal samples were collected at three time points and bacterial 16s rRNA genes were sequenced to profile the microbiota. As expected, cocaine-related behavior varied among subjects, such that a quartile split identified low and high responders on each measure, as well as an overall addiction index. Although beta diversity in the microbiota at baseline and after short access did not predict membership in high or low addiction quartiles, linear discriminant analysis (LDA) identified taxa that were more robustly represented in low or high responders. Beta diversity after long access was different among quartiles, as were several specific taxa. Investigation of baseline differences revealed that high relative abundance of *Akkermansia muciniphila* predicted future low response rates, whereas *Ruminococcaceae* predicted high response. This study is the first to report that microbiota variability reflects levels of cocaine intake and that microbial profiles might facilitate diagnosis and identify risk factors predictive of drug vulnerability.

**Significance Statement:** Microbial organisms inhabiting the gut of animals appear to influence organismal function through various signaling pathways, ultimately affecting behavior and disease vulnerability. This experiment investigates links between gut bacteria and vulnerability to addiction-related behaviors in adult male rats. Not only did gut bacterial profiles change as a result of cocaine intake but also gut bacterial profiles before any exposure to cocaine predicted which animals would be high or low addiction-prone individuals. These results suggest that microbial profiles might facilitate diagnosis and identify risk factors predictive of drug addiction.

## Introduction

An estimated 165 million Americans over the age of 12 used addictive substances in the past month (Welty et al., 2019), despite findings that drug addiction is associated with deleterious outcomes for many individuals and society on the whole (Volkow and Boyle, 2018). Notably, only about 15-30% of individuals who initiate drug use develop chronic use habits and addiction (Wang et al., 2012; Anthony et al., 1994; Saunders and Robinson, 2013; Franken et al., 2000), but the factors that underlie addiction vulnerability are still poorly understood. Effective treatments are also lacking (Fischer et al., 2015; Simpson et al., 1999). Thus, continued investigations of novel approaches to treat, predict, and prevent cycles of drug abuse and addiction are necessary.

The gut-brain axis is a bidirectional communication pathway between the gastrointestinal tract and the central nervous system (Sudo et al., 2004; Martin et al., 2018; Mayer et al., 2014; Cho and Blaser, 2012), and it appears to be involved in neuropsychiatric disorders (Kim and Shin, 2018; Cryan and Dinan, 2012; Diaz Heijtz et al., 2011), perhaps including substance use disorder (Kiraly et al., 2016; Meckel and Kiraly, 2019; Dinan and Cryan, 2017; Simpson et al., 2020). Microbes that inhabit the gut comprise the gut microbiota and maintain the local environment, carry out metabolic functions, and contribute to immune responsivity, among other critical roles. They produce molecules that signal locally, activate neuronal projections to the brain, and enter the bloodstream for distribution throughout the body (Cryan and Dinan, 2012; Cryan et al., 2019). Variability in microbial communities among subpopulations of humans and other animals may contribute to individual differences in general behavioral characteristics, disease vulnerability, and symptomatology, but the ability of microbial profiles to predict vulnerability to addiction has not been explored.

Both clinical studies and animal models suggest links between the gut microbiome and drugs of abuse. Microbial profiles differ among individuals abusing alcohol (Engen et al., 2015), opioids (Gicquelais et al., 2020), or cocaine (Volpe et al., 2014), compared with healthy volunteers, although which came first, the drug intake or the altered microbiome, is not clear. In rats, repeated exposure to volatilized cocaine significantly altered both diversity and richness in gut bacterial communities (Scorza et al., 2019). Moreover, gut bacterial depletion via non-absorbable oral antibiotics in mice is associated with elevations in cocaine conditioned place preference (CPP), perhaps related to reduced production of short-chain fatty acids (Kiraly et al., 2016). Methamphetamine CPP correlates with reduced short-chain fatty acid levels in rat fecal samples as well (Ning et al., 2017). Antibiotic-induced gut microbial depletion also alters the neuronal ensembles that are activated by oxycodone intoxication and withdrawal (Simpson et al., 2020). Finally, links between gut dysbiosis and anxiety, depression, and heightened stress responsivity in animal models suggest an indirect path through which gut dysbiosis could create vulnerability to drug reward and reinforcement (Bercik et al., 2010; Diaz Heijtz et al., 2011).

The present study tested the hypothesis that drug use and microbiota composition are linked in such way that microbiota profiles can predict susceptibility to drug use and that drug use can alter microbiota composition, to promote further drug seeking behavior, resulting in a vicious cycle of drug abuse. Specifically, we predicted that the gut microbial communities characterized through fecal sample analysis would be different between addiction-resistant and addiction-prone adult male rats, with addiction vulnerability defined by a battery of cocaine-related behavioral tests including self-administration, escalation, and compulsive drug-seeking in the presence of an electric shock stimulus.

## Materials and Methods

### Animals

Male rats of heterogenous stock (HS) were provided by Dr. Leah Solberg Woods (Medical College of Wisconsin, now at Wake Forest University School of Medicine). Rats were housed two per cage on a reverse 12 h/12 h light/dark cycle (lights off at 0800 h) in a temperature (20–22°C)- and humidity (45–55%)-controlled animal facility with ad libitum access to water and food. All experiments were designed to minimize animal suffering. All of the procedures were conducted in adherence to the National Institutes of Health Guide for the Care and Use of Laboratory Animals and were approved by the Institutional Animal Care and Use Committee at the [Author University].

### Drugs

Cocaine HCl (National Institute on Drug Abuse, Bethesda, MD) was dissolved in 0.9% saline (Hospira, Lake Forest, IL) at a dose of 0.5 mg/kg per 0.1 ml infusion and self-administered intravenously.

### Intravenous catheterization

The animals were anesthetized by isoflurane inhalation, and intravenous catheters were implanted in the right jugular vein using a modified version of a procedure that was described previously (Grimm et al., 2001; Doherty and Frantz, 2013). The catheter assembly consisted of an 18 cm length of Micro-Renathane tubing (0.023-inch inner diameter, 0.037-inch outer diameter; Braintree Scientific, Braintree, MA, USA) attached to a guide cannula (Plastics One, Roanoke, VA, USA), which was bent at a near right angle, embedded in dental acrylic, and anchored with mesh (2 cm square). The jugular vein was punctured with a 22-gauge needle, and then the tubing was inserted and secured inside the vein with suture thread. The catheter portal exited through an incision on the back and sealed with a plastic cap and metal cover. Catheters were flushed daily with heparinized saline (10 U/ml of heparin sodium; American Pharmaceutical Partners, Schaumburg, IL, USA) in 0.9% bacteriostatic sodium chloride (Hospira, Lake Forest, IL, USA) that contained 20 mg/0.2 ml of the antibiotic Cefazolin (Hospira, Lake Forest, IL, USA). After recovery from surgery, rats were tested in several behavioral assays, per the experimental timeline (**Figure 1**). All behavioral testing was conducted during the dark phase.

**Figure 1:**
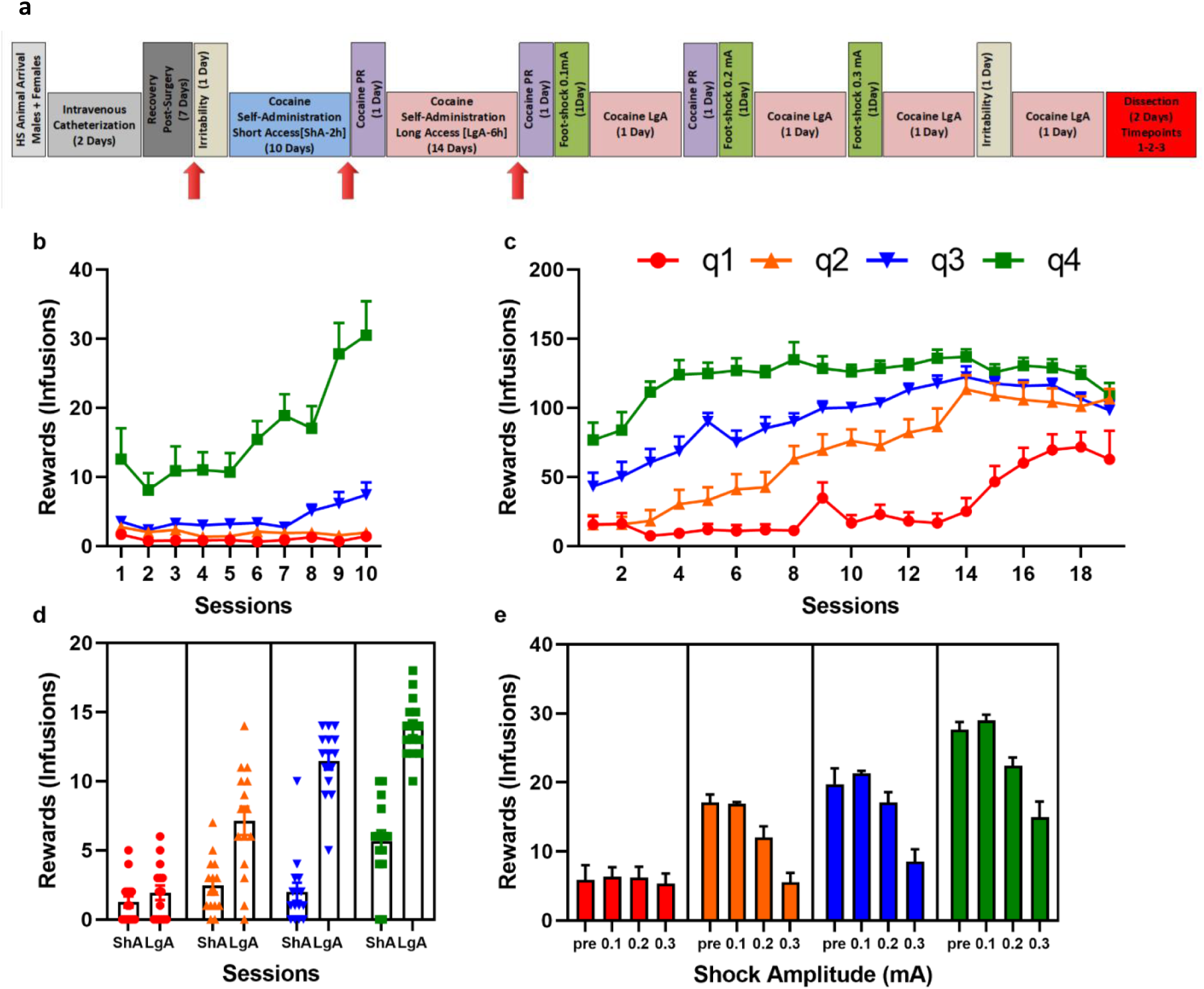
Behavioral testing timeline and behavioral outcomes. **a.**The experimental timeline indicates the order of procedures carried out with all subjects. Fecal matter was collected at the timepoints indicated by the arrows.Based on the number of cocaine infusions or behavior on the PR schedule, rats were divided intofour quartiles(q)(from low to high; green, yellow, orange, red). Panels show rewards earned undervarious conditions:**b.**shortaccess (2 h/day), **c.** long access (6 hr/day), **d.** Progressive ratio (PR)schedule of reinforcement, **e.**compulsive drug seeking in the face of footshock paired with cocaine infusions.

### Operant training

Self-administration was performed in operant chambers equipped with two retractable levers (Med Associates, St. Albans, VT, USA). Cocaine was delivered through an infusion pump that was activated by responses on the right lever (active), resulting in the delivery of cocaine (0.5 mg/kg per 0.1 ml). Responses on the left lever (inactive) were recorded but had no scheduled consequences. The rats were first trained to self-administer cocaine under a fixed-ratio 1 (FR1) schedule of reinforcement in daily 2-h, short-access (ShA) sessions for 10 days.

Presentation of cocaine reinforcers was paired with onset of a cue light over the active lever and followed by a 20-s timeout (TO), during which lever presses were counted but did not result in cocaine infusion. Subsequent 6-h, long access (LgA) self-administration sessions over 14 days allowed rats to escalate their cocaine intake (Ahmed and Koob, 1998).

### Progressive Ratio

At the end of the ShA and LgA phases, a progressive ratio (PR) schedule of reinforcement was used to assess the break point in lever-press responding, a measure of the reinforcing value of a reward (Stafford et al., 1998; Hodos, 1961). The PR response requirements necessary to receive a single drug dose increased according to this equation: [5e ^(injection numbers x 0.2)^] – 5, resulting in the following progression of response requirements: 1, 2, 4, 6, 9, 12, 15, 20, 25, 32, 40, 50, 62, 77, 95, 118, 145, 178, 219, 268, etc. The breakpoint was defined as the last ratio attained by the rat prior to a 60 min period during which a ratio was not completed. Progressive ratio testing was repeated two times after the LgA phase.

### Compulsivity

After LgA sessions and PR testing, rats were placed in the self-administration chamber for 1 h and tested for compulsive-like behavior. In this procedure, rats were allowed to self-administer cocaine on an FR1 schedule, but 30% of the reinforced responses were paired with a contingent footshock (0.1 mA, 0.5 s). In two more sessions spaced three days apart, the shock intensity increased to 0.2 mA and 0.3 mA successively.

### Feces collection

At times marked in **Figure 1a**, rats were held by the base of the tail so that the front paws were on a solid surface and the back paws were lifted off the surface. The back paws were gently moved up and down off of the surface until a fresh fecal pellet was released, collected immediately into a sterile tube, put on dry ice, then frozen at -80°C for storage until overnight shipping to Georgia State University’s Neuroscience Institute for storage at -80°C until processing.

### Fecal microbiota analysis by 16S rRNA gene sequencing

16S rRNA gene amplification and sequencing were done using the Illumina MiSeq technology following the protocol of Earth Microbiome Project with modifications (www.earthmicrobiome.org/emp-standard-protocols) (Gilbert et al., 2014; Caporaso et al., 2012). Bulk DNA was extracted from frozen feces using a Qiagen Power Fecal DNA Isolation Kit with mechanical disruption (bead-beating). The 16S rRNA genes, region V4, were PCR amplified from each sample using a composite forward primer and a reverse primer containing a unique 12-base barcode, designed using the Golay error-correcting scheme, which tagged PCR products from respective samples (Caporaso et al., 2012).We used the forward primer 515F 5’-*AATGATACGGCGACCACCGAGATCTACACGCT*XXXXXXXXXXXX**TATGGTAATT*G T***<u>GTGYCAGCMGCCGCGGTAA</u>-3’: the italicized sequence is the 5’ Illumina adapter, the 12 X sequence is the Golay barcode, the bold sequence is the primer pad, the italicized and bold sequence is the primer linker, and the underlined sequence is the conserved bacterial primer 515F. The reverse primer 806R used was 5’-*CAAGCAGAAGACGGCATACGAGAT***AGTCAGCCAG*CC*** <u>GGACTACNVGGGTWTCTAAT</u>-3’: the italicized sequence is the 3’ reverse complement sequence of Illumina adapter, the bold sequence is the primer pad, the italicized and bold sequence is the primer linker, and the underlined sequence is the conserved bacterial primer 806R. PCR reactions consisted of Hot Master PCR mix (Quantabio, Beverly, MA, USA), 0.2 mM of each primer, and 10-100 ng template, with reaction conditions over 3 min at 95°C, followed by 30 cycles of 45 s at 95°C, 60s at 50°C, and 90 s at 72°C on a Biorad thermocycler. PCR products were purified with Ampure magnetic purification beads (Agencourt, Brea, CA, USA) and visualized by gel electrophoresis. Products were then quantified (BIOTEK Fluorescence Spectrophotometer) using Quant-iT PicoGreen dsDNA assay. A master DNA pool was generated from the purified products in equimolar ratios. The pooled products were quantified using Quant-iT PicoGreen dsDNA assay and then sequenced using an Illumina MiSeq sequencer (paired-end reads, 2 x 250 bp) at Cornell University, Ithaca.

### Genetic sequence analysis

16S rRNA gene sequence analysis was performed as previously described (Chassaing et al., 2015). Sequences were demultiplexed, quality-filtered using the Quantitative Insights Into Microbial Ecology (QIIME) software package (Caporaso et al., 2012), and forward and reverse Illumina reads were joined using the fastq-join method (http://code.google.com/p/ea-utils) (Aronesty, 2013). QIIME default parameters were used for quality filtering and sequences were assigned to operational taxonomic units (OTUs) using the UCLUST algorithm (Edgar, 2010), with a 97% threshold of pairwise identity, and classified taxonomically using the Greengenes reference database (McDonald et al., 2012). Principal coordinate analysis of the unweighted UniFrac distances was used to assess the variation between samples (beta diversity) on rarefied OTU tables (Lozupone et al., 2011). Alpha diversity and rarefaction curves were calculated and displayed using observed OTUs as the variable of interest on rarefied OTU tables.

### Identification of bacteria predicting drug use

Predictive analysis was conducted by generating Receiver Operating Characteristic (ROC) curves, and area under the curve (AUC), as adapted from (Sokol et al., 2019) using multiple logistical regression in PRISM 8.3.0 Graphpad (v 2019). In brief, regression plots of the species relative abundance was plotted against bacteria present in sample (binary: yes/no). Artifacts due to phylogeny differences were discarded such that curves represent lowest similar taxonomic identification. ROC curve quality was assessed based on previous standards (Mandrekar, 2010).

### Statistics

Statistics on behavioral outcomes, including median splits and z-score calculations were conducted with IBM SPSS v. 23, and corresponding graphs were generated using PRISM 7 GraphPad (San Diego, California, USA). Generation of robust bacterial differences between groups, using Linear discriminant analysis Effect Size (LEfSe) was accomplished using the Galaxy tool (http://huttenhower.sph.harvard.edu/galaxy/) (Goecks et al., 2010). Galaxy was also used to generate Linear Discriminant Analysis (LDA) bar charts and corresponding cladograms. Determination of predictive biomarkers was accomplished via a combination of LEfSe baseline output data, highly abundant taxonomic groups observed in the present samples, and the literature on bacterial groups previously associated with conditions related to substance use disorder. Diversity plots and relative abundance stacked bar charts were generated via QIIME. Differences in distinct clustering in PCA plots was assessed via PERMANOVA method using vegan R-package through QIIME.

## Results

### Identification of low- and high-vulnerability subpopulations based on behavioral outcomes

Rats self-administered cocaine under four different experimental conditions, per the timeline in **Figure 1a**. In each condition, variability in response rates allowed division of subjects into low vs. high responder subgroups, using a quartile split separately for each measure (**Figure 1b-e**). Thus, all the behavioral variables were analyzed using a separate quartile analysis with Q1 capturing the lowest 25% of the population and Q4 capturing the highest 25% of the population for the given dependent variable. During ShA self-administration (2 h/day), the number of cocaine infusions (rewards) earned per session was between 0 and 90, with an overall average of 6.13 +/- 6.17 (Fig. 1b). The bottom 25% of animals based on the sum of infusions over all ShA sessions were considered to be low responders (or addiction resistant) and had an average of 0.97 +/- 0.1 rewards earned per session. The top 25% of rats, labeled as high responders (or addiction vulnerable), had an average of 16.3 +/- 1.21 per session. During LgA escalation conditions (6 h/day), cocaine infusions ranged from 0 to 277 with an overall average of 77.14 +/- 1.59. Low responders based on the sum of all infusions during 14 LgA sessions averaged 27.13 +/- 2.15 infusions per session, whereas high responders averaged 121.8 +/- 2.05 infusions per session (**Fig. 1c**). During PR testing, the total number of infusions earned appeared higher when tested after LgA escalation sessions compared with ShA sessions in the three upper quartiles but showed more overlap across testing points in the low responders (**Figure 1d**). Quartiles were defined by the sum of infusions earned over all three PR sessions, whereas only the first two PR sessions are plotted. Rats in the lowest quartile averaged just 1.6 +/- 0.33 infusions, requiring 3.47 +/- 0.68 active lever presses during PR, whereas rats in the highest quartile earned an average of 9.77 +/- 0.89 infusions with 49.7 +/- 7.39 active lever presses during PR after LgA. Under intermittent shock conditions, most rats earned fewer cocaine infusions as the shock intensity increased, although the low responder quartile earned low numbers of infusions regardless of shock intensity (**Figure 1e**), with quartiles defined based on the sum of infusions over all four shock values. To summarize these data mathematically, the sum of infusions earned by each rat on each measure (LgA, PR, and shock) was transformed into a Z-score and also averaged to create an overall Addiction Index (**Figure 2**). The lowest 25% of responders (Q1; addiction resistant) and the top 25% of responders (Q4; addiction vulnerable) on each individual behavior were used to categorize rats for comparisons of microbial profiles, as were the lowest vs. highest quartiles as defined by the overall addiction index. Notably, the mid-range quartiles were not included in microbial comparisons, except for the cohort comparison (**Supplemental Figure 1**).

**Figure 2:**
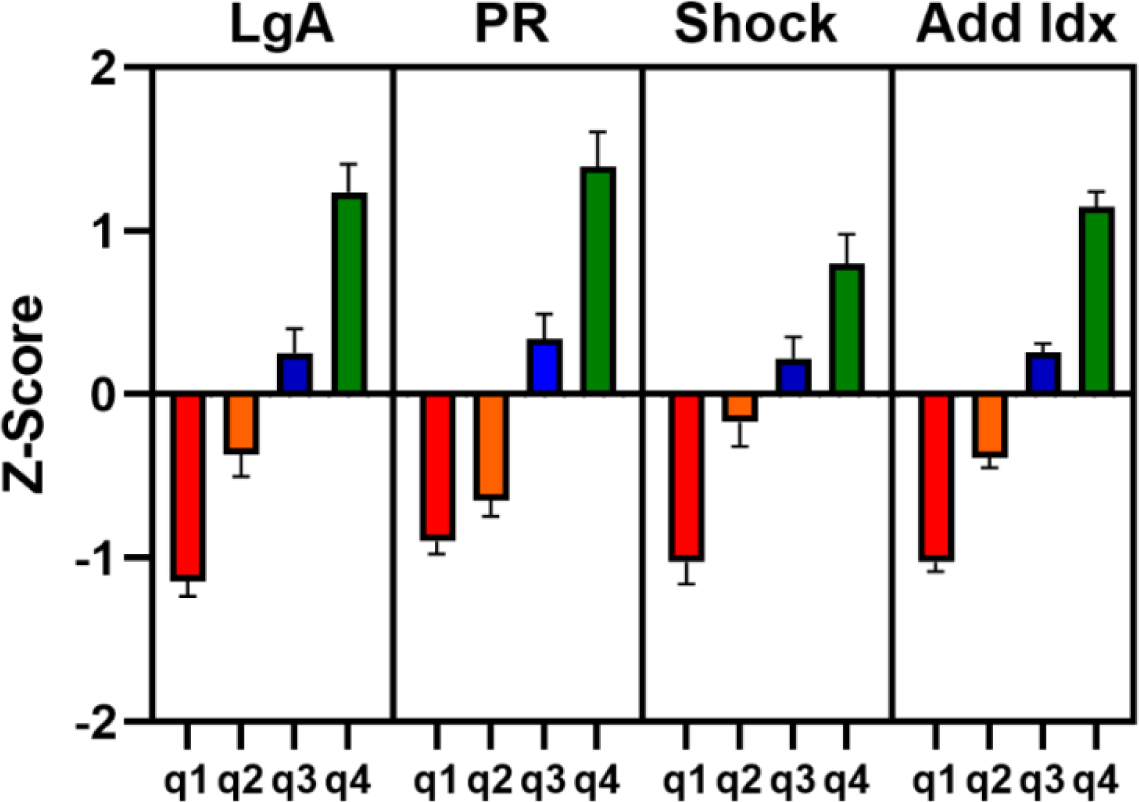
Cocaine-related behavioral outcomes categorized into quartiles by z-scores. The left three panels show individual z-score distributions for low and high responders on the long-access, PR, and compulsive (shock) conditions. The right panel shows an average of scoring on all three measures, creating an overall addiction index and plotting z-score distributions for low and high responders. All subjects self-administered cocaine under all conditions.

### Short-access cocaine self-administration induced modest alteration in the intestinal microbiota composition

To assess the impact of ShA cocaine conditions on the intestinal microbiota, we used the addiction index z-scores to divide subjects into low (Q1) and high (Q4) vulnerability groups and compared the microbiota composition from fecal samples taken after the last ShA session. Overall microbial communities did not cluster into separate populations, i.e. beta diversity was not significantly different based on principal coordinates analysis (**Figure 3a**). Yet individual bacterial groups did vary in relative abundance, as shown in the LEfSe cladogram (**Figure 3b**), bar chart (**Supplemental Figure 2b**), and **Table 1**. Specifically, the order *Enterobacteriales*, family *Enterobacteriaceae*, and genus *Anaerostipes*, in the class *Gammaproteobacteria*, were more robustly expressed in the low responders vs the high responders at this time point, although none of these groups was found in particularly high levels in either addiction group (data not shown). Based on observed OTUs, no differences in alpha diversity after short access cocaine self-administration were seen in low vs high responders. (**Supplemental Figure 2a**).

**Figure 3:**
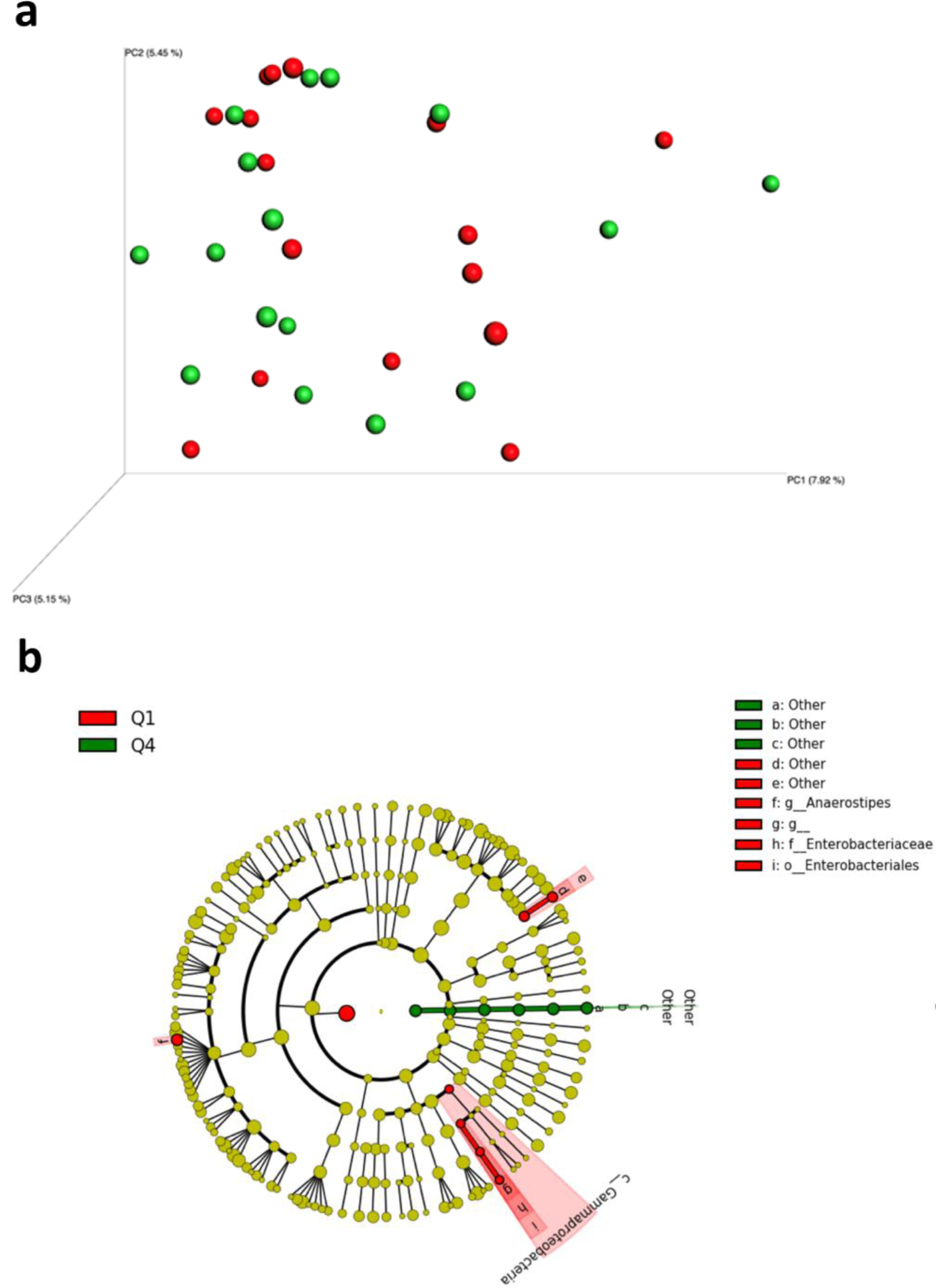
Impact of short-access cocaine self-administration on microbiota composition. **a.** Principal coordinates analysis (PCoA) of the unweighted UniFrac distance matrix of the bottom quartile (Q) of cocaine resistant rats on the overall addiction index (Q1, red) and top quartile of cocaine vulnerable animals (Q4, green), shows no significant clustering based on sequencing 16s rRNA bacterial genes from fecal samples collected after short-access conditions. **b.** Taxonomic cladogram obtained from linear discriminant analysis effect size (LEfSe) highlights specific bacterial taxa that were relatively more abundant in Q1 (red) or more abundant in Q4 (green). Minimum LEfSe score was 2.0.

**Table 1.**
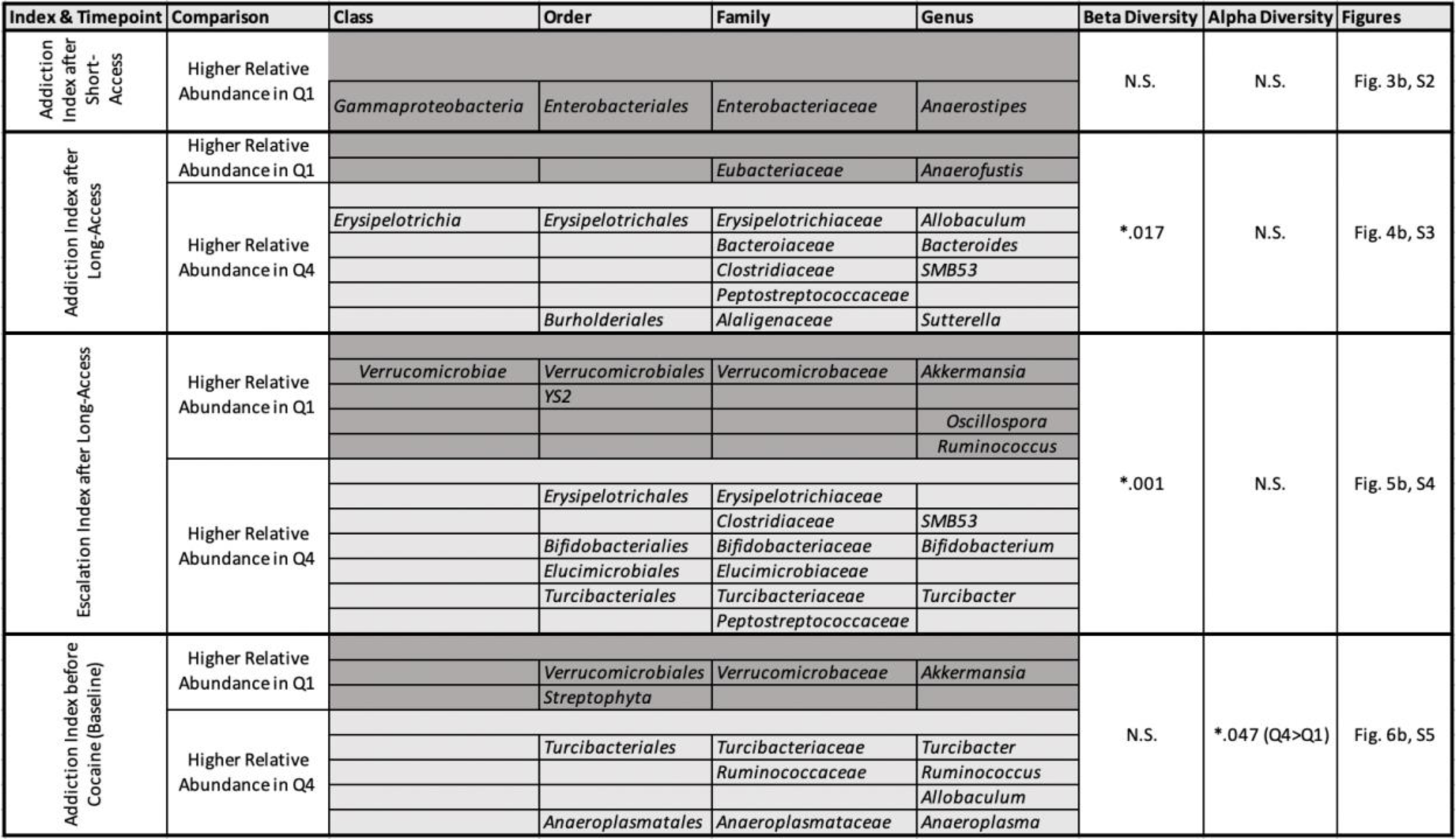
Summary of differential bacterial abundance in fecal samples.

### Long-access cocaine self-administration induced gut microbiota dysbiosis

To assess the impact of longer (6 hr/day) cocaine self-administration sessions on the intestinal microbiota, we used the addiction index z-scores to divide subjects into low (Q1) and high (Q4) responder quartiles and compared the microbiota composition from fecal samples taken after the last of 14-straight LgA sessions. Microbial communities were clustered into distinct populations, as determined by PCoA of the unweighted UniFrac distance matrix (**Figure 4a**). Moreover, individual bacterial groups varied in relative abundance, as demonstrated with LEfSe results plotted in a cladogram (**Figure 4b**), bar chart (**Supplemental Figure 3**), and Table 1. Specifically, the class *Erysipeltotrichi*, order *Erysipetotrichales*, family *Erysipelotrichaceae,* and genus *Allobacullum* were more robustly expressed in the high responders vs low responders at this time point. *Bacteroidaceae* and *Bacteroides*, as well as *Clostridiaceae* and *SMB53*, *Peptostreptococcaceae*, and the *Burholderiales*, *Alaligenaceae*, and *Sutterella* clade were also more robust in the high responders. On the other hand, drug resistant animals showed relatively higher abundance of the family *Eubacteriacease* and genus *Anaerofustis*. Among these, only the phylum *Bacteroidetes* was identified at high proportions in the samples (**Supplemental Figure 3b**). Based on Observed OTUs, no differences in alpha diversity after short access cocaine self-administration were seen in low vs high responders (**Supplemental Figure 3a**).

**Figure 4:**
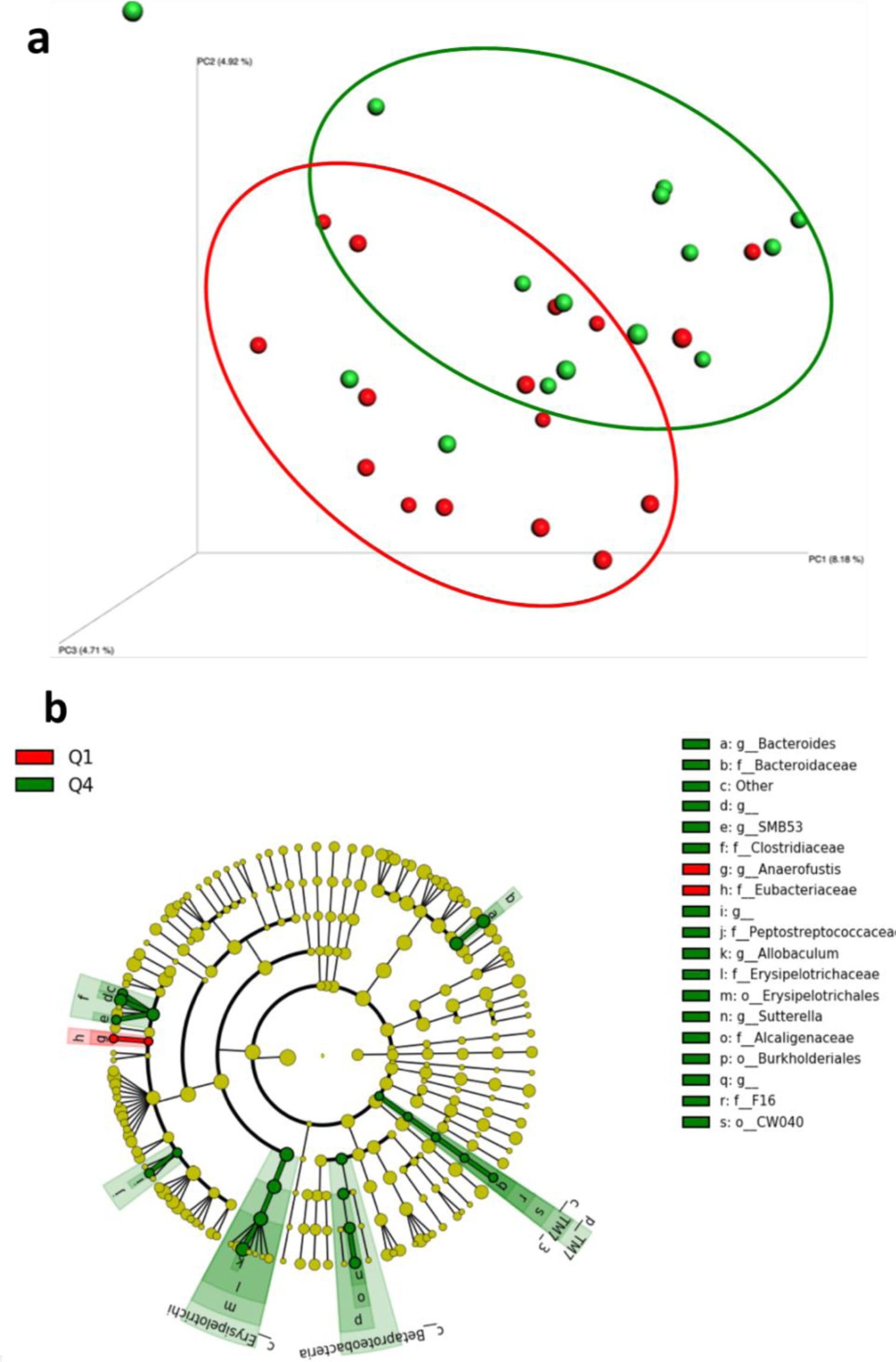
Impact of long-access cocaine self-administration on microbiota composition. **a.** Principal coordinates analysis (PCoA) of the unweighted UniFrac distance matrix of the lowest quartile of responders defined by the overall addiction index (Q1, red) and highest quartile of responders (Q4, green), shows differential clustering based on sequencing 16s rRNA bacterial genes from fecal samples collected after these long-access conditions (p=0.017). **b.** Taxonomic cladogram obtained from LEfSe highlights specific taxa that were relatively more abundant in Q1 (red) or Q4 (green). Minimum LEfSe score was 2.0.

In addition to using the addiction index z-scores, we also used the long-access (escalation) z-scores to subdivide animals to consider the impact of long-access cocaine self-administration on microbial profiles. With this differentiation, we again observed a significant difference in beta diversity between high and low responders (**Figure 5a**). We also observed overlap in bacterial subpopulations showing a greater relative abundance in high responders: the order *Erysipelotrichales* and family *Erysipilotrichaceae*, as well as family *Clostridiaceae* and genus *SMB53* (**Figure 5b**, **Supplemental Figure 4b, Table 1**). In addition, several groups showed higher relative abundance in high responders defined by this escalation z-score but not the addiction index z-score: orders *Bifidobacteriales*, *Elusimicriobiales*, and *Turicibacterales*, families *Bifidobacteriaceae*, *Elusimicrobiaceae*, *Turicibacteraceae*, and *Peptostreptococcaceae*, and genera *Bifidobacterium* and *Turcibacter*. Conversely, several groups showed higher relative abundance in the low responders: class *Verrucomicrobiae*, order *Verrucomicrobiales*, family *Verrucomicrobiaceae*, and genus *Akkermansia*, as well as order *YS 2* and genera *Oscillospira* and *Ruminococcus*. Using the escalation index as the determining factor for cocaine vulnerability resulted in no alpha diversity differences, as defined by observed OTUs (**Supplemental Figure 4a**). The relative abundance charts compare the gut bacteria phenotypes between high and low responders at the phylum level (**Supplemental Figure 4c**).

**Figure 5:**
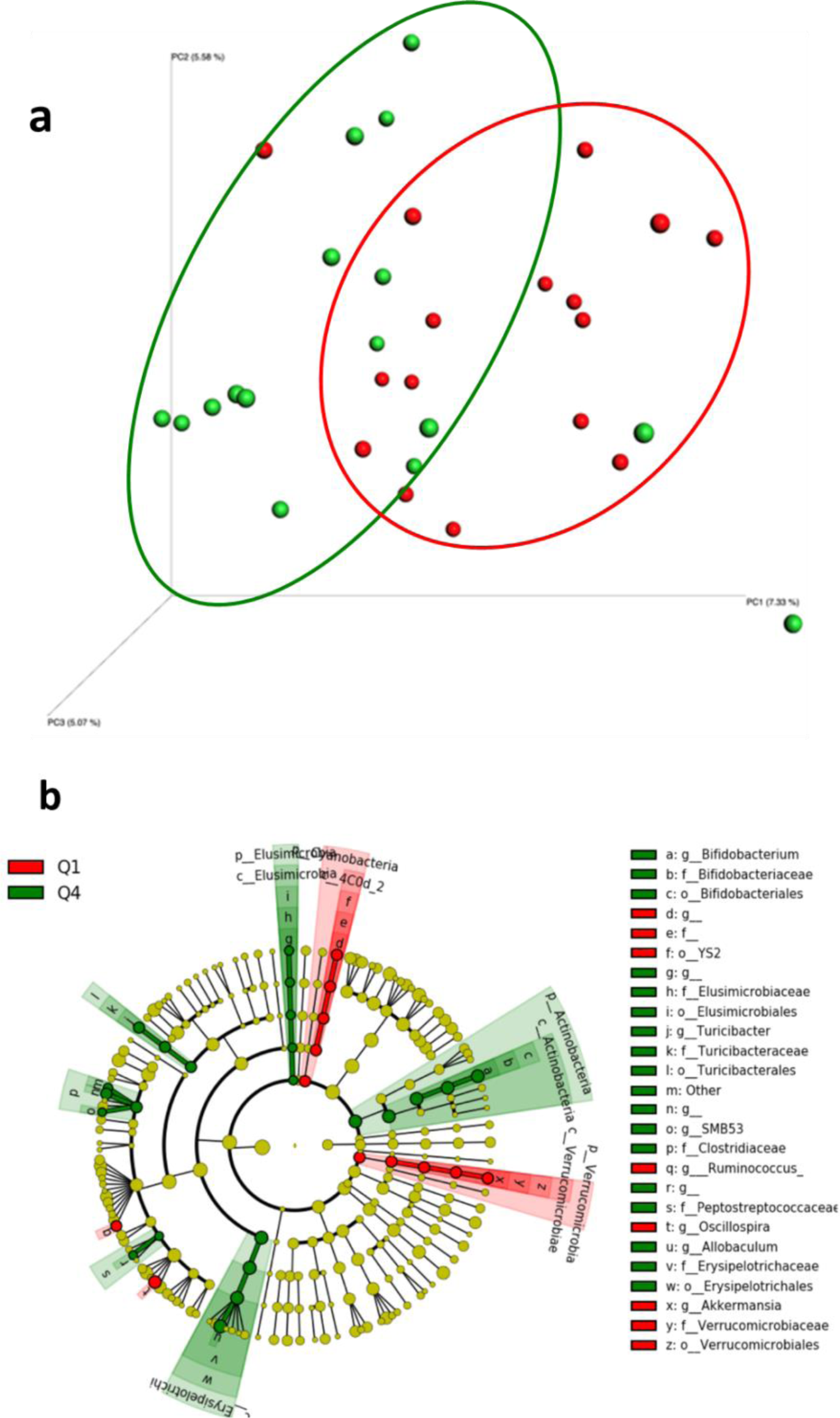
Impact of long-access cocaine self-administration on microbiota composition, with subpopulations defined by long-access (escalation) z-score. **a.** Principal coordinates analysis (PCoA) of the unweighted UniFrac distance matrix of the bottom quartile (Q) of cocaine resistant rats on the specific escalation z-score (Q1, red) and top quartile of cocaine vulnerable animals (Q4, green), shows differential clustering based on sequencing 16s rRNA bacterial genes from fecal samples collected after these long-access conditions (p=0.001). **b**. Taxonomic cladogram obtained from LEfSe highlights specific taxa that were relatively more abundant in Q1 (red) or Q4 (green). Minimum LEfSe score was 2.0.

### Microbial composition may predict future cocaine use

As noted in the timeline (**Figure 1a**), fecal samples were also collected before any exposure to cocaine. Microbiota composition analysis in these samples demonstrated that before cocaine exposure, microbiota profiles did not cluster differentially based on the likelihood of cocaine use later in the experiment, as presented in **Figure 6** when using the Addiction Index z-score. However, when looking at individual taxa contributions to future phenotypes via LeFSe analysis (**Supplemental Figure 5b**) and taxa summarization (**Supplemental Figure 5c**), we observed that certain bacterial groups were significantly different in future low vs high cocaine responders (**Table 1**). Specifically, the order *Verrucomicrobiales*, family V*errucomicrobaceae*, and species *Akkermansia muciniphila* had higher relative abundance in future low responders than high responders, as did the order *Streptophyta*. Conversely, the orders *Anaeroplasmatales* and *Turcibacterales*, families *Turicbacteracae*, *Anaeroplasmataceae*, and *Ruminococcaceae*, along with genera *Turicbacter, Aneroplasma*, *Ruminococcus*, and *Allobaculum* were all more robustly expressed in future high responders vs future low responders (**Figure 6** **and Supplemental Figure 5b**). Some of these clades belong to phyla that are among the most highly abundant in both low and high responders (*Actinobacteria, Firmicutes, Tenericutes,* and *Verrucomicrobia*; **Supplemental Figure 5c**). Moreover, future high responders had higher alpha diversity than future low responders, based on significant differences in the number of observed OTUs (p=0.047; **Supplemental Figure 5a**).

**Figure 6:**
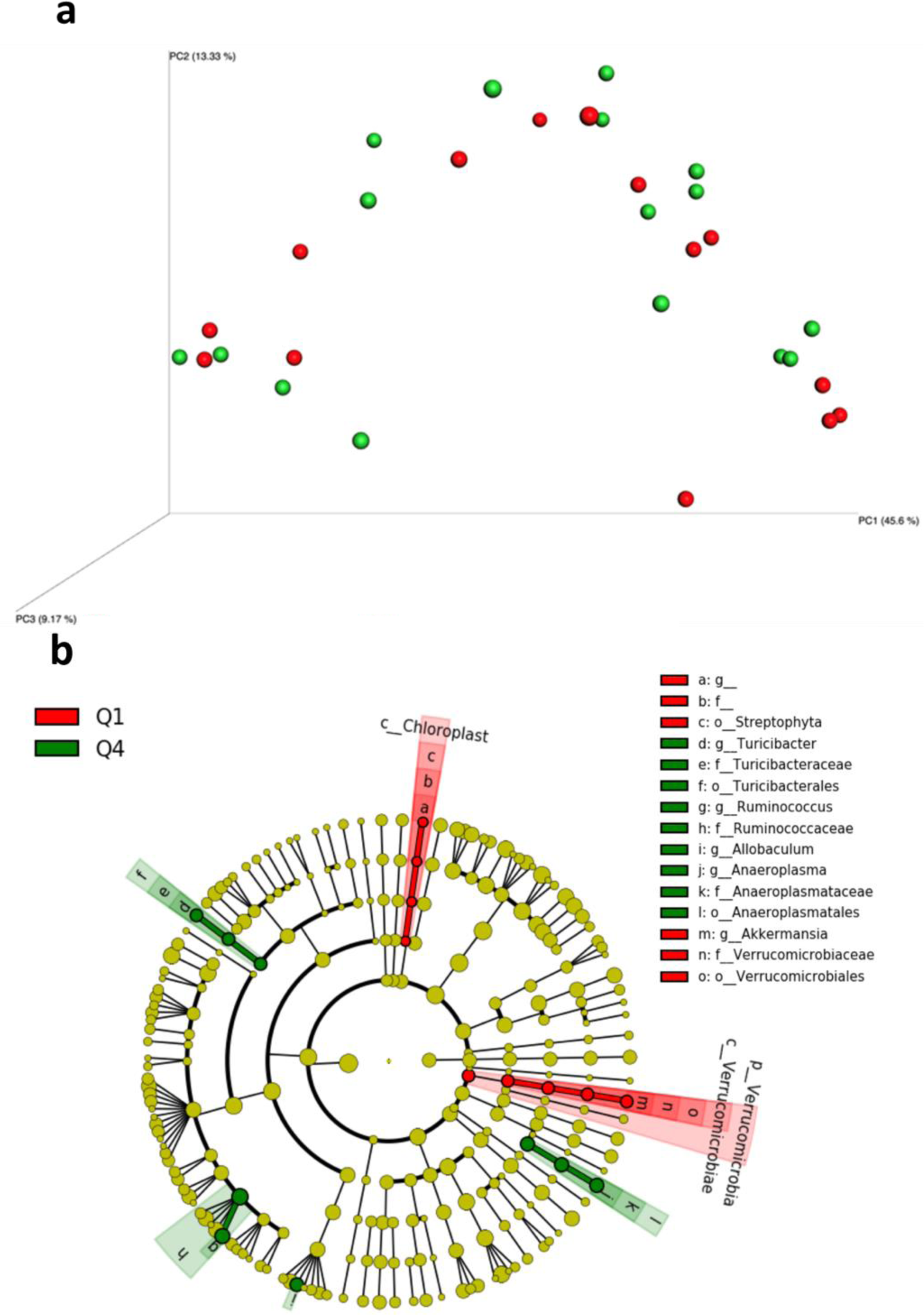
Using the microbiome to predict future cocaine sensitivity. **a.** Principal coordinates analysis (PCoA) of the unweighted UniFrac distance matrix of the bottom quartile (Q) of cocaine resistant rats defined by the addiction index (Q1, red) and top quartile of cocaine vulnerable animals on the addiction index (Q4, green) did not show differential clustering based on 16s rRNA bacterial genes from fecal samples collected before cocaine intake (baseline samples). **b.** Taxonomic cladogram obtained from LEfSe shows differential profiles between rats that go on to show low vs. high responsivity, with taxa relatively more abundant in Q1 resistant (red) or Q4 vulnerable (green) subpopulations, based on samples collected before cocaine. Minimum LEfSe score was 2.0.

Using output from the LEfSe, we chose several clades to test as potential biomarkers that might discriminate future membership in the low vs. high responder quartiles, based on the overall addiction index. The ROC curve plotting rates of true positive (sensitivity) against false positive (specificity) revealed that higher abundance of the species *Akkermansia muciniphila* showed excellent predictive value for membership in the low responder group (AUC=.8103), whereas the family *Ruminococcaceae* predicted high responsivity (AUC=0.7388; **Figure 7**).

**Figure 7:**
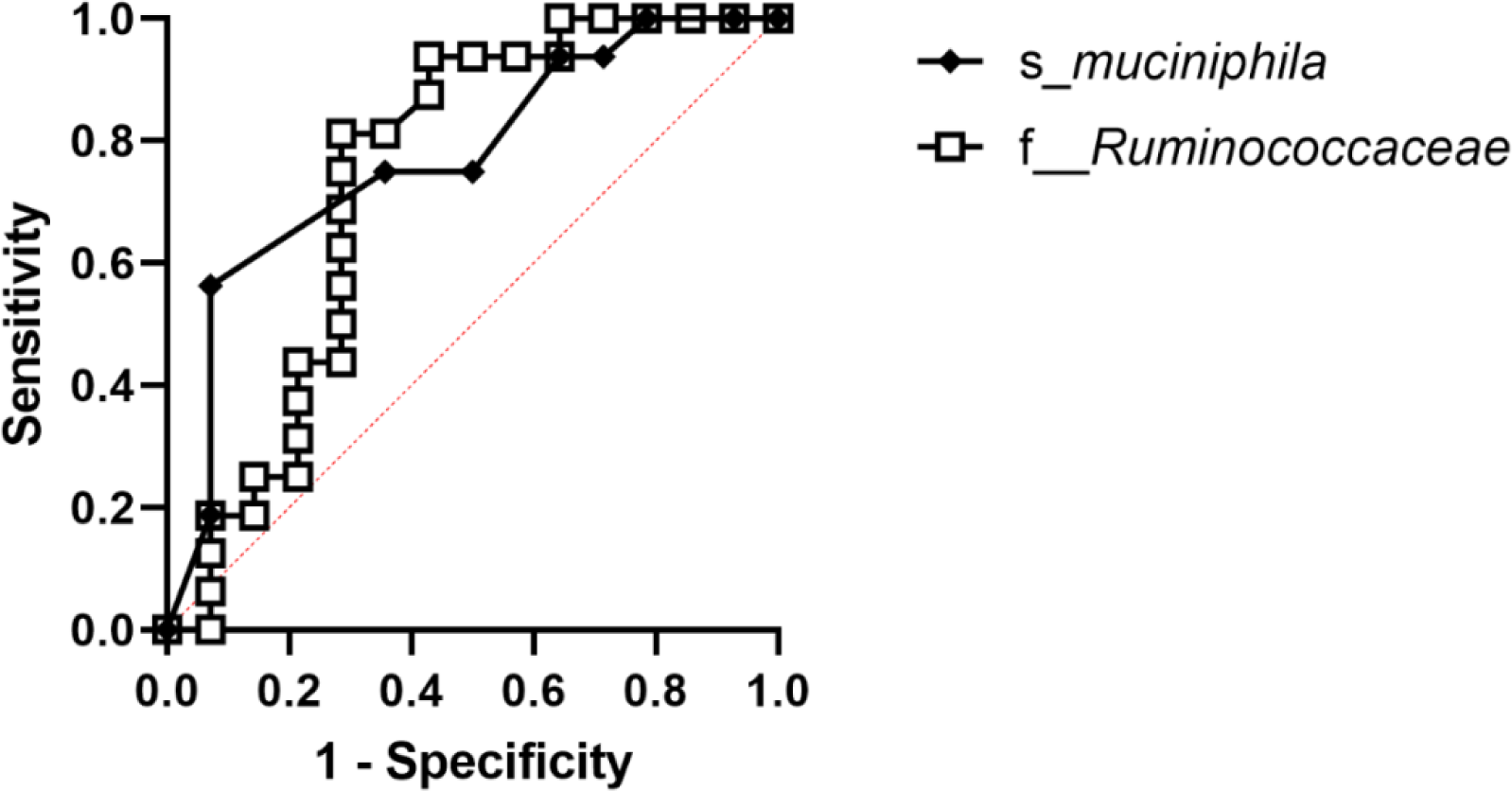
Specific taxa predict future cocaine sensitivity. Specific microbial signatures were analyzed from fecal samples obtained at baseline before any cocaine exposure, using a random forest algorithm to predict future phenotype contribution. The species *Akkermansia muciniphila* (AUC=0.8103) and family *Ruminococcaceae* (AUC=0.7388) both discriminated membership in low vs high future addiction index phenotypes. AUC, area under the curve.

Based on relevant literature linking the phyla *Firmicutes* and *Bacteroidetes* (or the ratio of their relative abundance) with obesity (John and Mullin, 2016), age (Mariat et al., 2009), and alcohol use disorder (Engen et al., 2015), we also plotted ROC curves for these clades. *Firmicutes* and *Bacteroidetes* separately showed poor discrimination of future cocaine sensitivity with AUCs of 0.6429 and 0.6027 (**Supplemental Figure 6a and b**), respectively. Similarly, the ratio of the two yielded poor predictive results (AUC=0.625; **Supplemental Figure 6c**).

Evidence suggests that the genus *Lactobacillus*, part of the order *Lactobacillaceae*, is important in psychological and social health and well-being. In rodents, probiotics containing *Lactobacillus* alleviate responsivity to moderate psychological stress (Messaoudi et al., 2010; Messaoudi et al., 2011) and reverse deleterious effects of psychological stress on the brain (Ait-Belgnaoui et al., 2013). Given evidence that addiction phenotypes are comorbid with neuropsychiatric conditions such as exaggerated stress responsivity (Regier, 1990), we sought to examine whether *Lactobacillus*-related bacterial populations at baseline could predict future drug use. ROC curves revealed that the order *Lactobacillaceae* showed fair discrimination (AUC=0.6897; **Supplemental Figure 7a**), while the genus *Lactobacillus* was a poor predictor (AUC=0.6272; **Supplemental Figure 7b**).

### Microbial profiles vary by experimental cohort

The behavioral assays carried out in this study required rats to be tested in cohorts of approximately 20 animals each. We therefore explored the impact of cohort on microbial profiles and noted significant differences in beta diversity (**Supplemental Figure 1**).

## Discussion

Cocaine use is a debilitating condition estimated to affect over 20 million people worldwide (Pomara et al., 2012). Despite the societal strain caused by the use of cocaine and other illicit substances, effective therapeutics have yet to be developed, in part due to the high variability in levels and patterns of intake (Davidson et al., 1993), a phenomenon mirrored in rodent models (Panlilio et al., 2003). Based on relatively new discoveries linking cocaine-related behavior to the gut microbiome (Kiraly et al., 2016; Meckel and Kiraly, 2019; Cryan and Dinan, 2012; Chivero et al., 2019), we tested the hypothesis that levels of cocaine responsivity would correlate with microbiota profiles in adult male rats, such that relative abundance of certain bacterial populations not only reflect but also predict vulnerability to cocaine reward and reinforcement. Thus, we investigated whether the microbiota could be a useful diagnostic tool to predict the severity of future cocaine use. The results supported the hypothesis that gut microbial communities in low vs high addiction-prone rats were different in ways that reflected the acute and long-term impact of cocaine intake on bacterial populations, and they also predicted subsequent sensitivity, revealing potential biomarkers for cocaine resilience or sensitivity.

The present behavioral and microbial assays were previously validated and are translationally relevant to addiction-like behaviors and established profiles of microbial composition. With regard to the behavioral assays measuring different aspects of cocaine reinforcement, fixed ratio schedules of reinforcement under short-access followed by long-access sessions may model escalation of drug intake with increased access to the drug (Ahmed and Koob, 1998); PR schedules may reflect motivation to obtain the drug (Stafford et al., 1998), and continued self-administration despite pairing of drug infusions with electric shock attempts to model the compulsive drug-seeking components of substance use disorder (Johnson and Kenny, 2010). Raw values and variability in cocaine-related behaviors were consistent with prior reports (Briand et al., 2008; George et al., 2008; Verheij et al., 2018), supporting the reliability and validity of the present outcomes. Quartile splits in behavioral response levels were used successfully to create low- and high-responding subgroups for each of these measures, along with a total addiction index that may reflect the multidimensional aspects of addiction, thereby elevating the relevance of the present analysis to the human condition. Microbiome profiling revealed levels of beta diversity and alpha diversity, as well as dominant bacterial taxa, that are similar to those reported in other studies of rodents (Li et al., 2017).

The impact of relatively low-level cocaine intake was assessed using fecal samples collected after the short-access phase of cocaine self-administration. Although the overall microbiota diversity and richness at this time point were not altered between animals that fell into the lowest vs highest quartiles of the overall addiction index, some interesting individual differences in relative abundance suggest the possibility that bacterial communities in low responders contained higher levels of some beneficial bacterial taxa. For example, *Anaerostipes* is among the many bacterial taxa known to produce short-chain fatty acids (Schwiertz et al., 2002; Duncan et al., 2004). Its higher abundance in low responders could support conclusions from prior reports that exogenous SCFA administration reduces cocaine’s reward value in mice (Kiraly et al., 2016). The family *Enterobacteriaceae* is abundant in the gut of newborn and juvenile humans (Bokulich et al., 2016; Yassour et al., 2016), perhaps suggesting a role in growth and development, while the overall class of *Gammaproteobacteria*, to which *Enterobacteriaceae* belong, is elevated after gastric surgery in both humans (Aron-Wisnewsky and Clement, 2014; Furet et al., 2010) and rodents (Liou et al., 2013), suggesting a role in recovery. Both these taxa were also more abundant in cocaine-resistant than cocaine-vulnerable animals in the present study.

In contrast to results seen after short-access cocaine self-administration, more robust differences in gut bacterial communities between low and high responding animals were observed after the long-access phase of self-administration, whether the quartile splits were based on LgA behavior alone or the overall addiction index. Beta diversity was significantly different in each case, suggesting that high cocaine intake during LgA sessions creates a distinct microbial environment in the gut. The two taxa described above as overrepresented in low vs high responders after short-access cocaine were no longer elevated after long-access. Yet *Anaerofustis* was, and it is also a SCFA producer of the family *Eubacteriaceae* in the class *Clostridia*. When categorizing rats based on the escalation index alone, two additional clades from the class *Clostridia* were higher among low responders: *Oscillospira*, which is associated with gastric dysfunction (Lam et al., 2012) and *Ruminococcus gnavus*, which is associated with Crohn’s disease and gut inflammation (Henke et al., 2019). In humans, several *Clostridia* species were also higher among non-opioid users, compared with opioid abuse patients (Acharya et al., 2017). Perhaps most notably, *Akkermansia muciniphila* was also higher among low responders at this time point. This species is well known for its inverse relationship with cardiometabolic disease and gut inflammation (Cani and de Vos, 2017; Naito et al., 2018). It is associated with integrity of the intestinal mucosal barrier, immune system activation, and production of SCFAs (van Passel et al., 2011). In clinical contexts, *A. muciniphila* has high relative abundance among non-obese, healthy controls (Xu et al., 2020; Dao et al., 2016) and can confer neuroprotection to mice similar to a ketogenic diet (Olson et al., 2018). On the other hand, chronic alcohol consumption lowers relative abundance of *A. muciniphila* whereas administration of *A. muciniphila* to patients with alcoholic liver disease ameliorates their symptoms (Grander et al., 2018).

Regardless of whether the overall addiction index or the escalation index was used to subdivide rats after LgA cocaine self-administration, the highest quartile of cocaine responders showed higher relative abundance of two different clades within the class *Clostridia: SMB53* and *Peptostreptococcaceae*, along with the genus *Allobaculum* from the class *Erysipelotrichia*. Interestingly, both methamphetamine and cocaine intake is related to higher relative abundance of *Allobaculum*, compared with healthy controls (Kosten, 2021; Scorza et al., 2019). A different set of clades differentiated high responders only when using the addiction index to separate vulnerability levels: *F16*, *Alcaligenaceae*, *Sutterella*, and *Bacteroides*. Consistent with this observation is the report that higher relative abundance of taxa in the phylum *Bacteriodetes* (which includes *Bacteroides*) was associated with cocaine use in humans (Volpe et al., 2014) and is higher among rodents given chronic alcohol (Yan and Schnabl, 2012). Moving on to those clades associated with high responders when defined by the escalation index instead, three taxa might be explored in the future for their specific relationship with higher doses of cocaine intake: *Elusicrobiaceae, Turcibacter*, and *Bifidobacterium*. Overall, the impact of these changes in the gut depends on the specific functionality provided by the clades independently, as well as their influence on one another in the gut. Furthermore, whether or not these differences in bacterial populations emerged as a consequence of cocaine intake or served to limit or drive cocaine intake cannot be known from this aspect of the present study. One approach to beginning to understand these distinctions is to investigate which clades showed differential relative abundance levels at baseline time points before cocaine intake.

Among the most promising translational components of this study is the potential to identify biomarkers of cocaine vulnerability. It is possible that the bacterial taxa that were higher at baseline among future low responders could be protective, whereas the taxa that were higher among future high responders could be associated with vulnerability. The most interesting of these might be *Akkermansia muciniphila*, which was higher at baseline among future low responders. Not only that, but it also remained higher among low responders throughout long-access testing. Further statistical exploration using the ROC curve suggested this species as an excellent predictor of belonging in the low addiction-prone subpopulation.

Conversely, perhaps playing a role in vulnerability are the *Allobaculum* and *Turcibacter* populations that were higher among high responders at baseline and after long-access testing. The genus *Anaeroplasma* was also higher at baseline among future high responders, warranting further investigation. Interestingly, the genus *Ruminococcus* was higher in high responders at baseline and was a strong predictor of membership in the high addiction-prone subpopulation, according to the ROC analysis. Nevertheless, the *Ruminococcus gnavus* species, per se, was higher in low responders later in the experiment, suggesting that its decline relative to other taxa could be an important factor in the behavioral switch to high response levels. Indeed this bacterial family has been associated with drug reward, impulsivity, attention, and locomotor activation in animal models (Ning et al., 2017; Peterson et al., 2020), as well as obesity and Alzheimer’s Disease in humans (Crescenzo et al., 2017; Zhuang et al., 2018; Ticinesi et al., 2018). On the other hand, though, *Ruminococcus*, a SCFA producer (Jiang et al., 2015; Acharya et al., 2017), appears lower in opioid users compared to healthy controls (Acharya et al., 2017), higher in abundance during abstinence from alcohol (Stärkel and Schnabl, 2016), and was inversely related to fibrosis in non-obese patients (Lee et al., 2020), leaving the role of *Ruminococcus* unclear. The present ROC curve analysis results did not support the use of *Lactobacillus* as a predictor of future membership in the low or high vulnerability quartiles, nor *Firmicutes*, *Bacteroidetes*, nor the *Firmicutes/Bacteroidetes* ratio.

The mechanisms through which cocaine alters the gut microbiome remain to be determined. General perturbations to gut function have long been associated with cocaine use and abuse (Brown et al., 1994; Gibbons et al., 2009). Cocaine exposure in mice triggers pro-inflammatory responses, compromises the mucosal lining of the gut, and alters microbiota composition (Chivero et al., 2019). As a sympathomimetic drug, cocaine increases gastric motility, thereby completely changing the timescale over which gut metabolism can be carried out. It also activates alpha-adrenergic receptors in the mesentery, leading to gastric ulcers and perforations in the gut membrane (Gourgoutis and Das, 1994; Cregler and Mark, 1986; Gibbons et al., 2009). Cocaine also reduces bloodflow to the gut, disrupts healthy diet, and hinders exercise patterns (Gibbons et al., 2009; Cregler, 1989), each of which also affects the gut microbiota indirectly (Luna and Foster, 2015; Maslowski and Mackay, 2011; Sandhu et al., 2017; Kang et al., 2014; Allen et al., 2015; Tang et al., 2017). These changes in the gut milieu are likely to favor certain bacterial populations while inhibiting the growth of others.

Although this is among the first reports to characterize the impact of cocaine on the gut microbiota while also suggesting diagnostic tools to predict the likelihood of drug vulnerability, two limitations are noted. First, microbial profiling from fecal samples rather than colon tissue restricts the view of the actual microbial environment in the gut (Stanley et al., 2015). The benefits of non-invasive, repeatable fecal sampling clearly outweigh this limitation, however. Second, the present subjects were tested in three cohorts, which showed baseline differences in the microbiome, thereby calling into question the generalizability of the identification of specific bacterial correlates with behavioral outcomes. This observation likely mirrors the human condition with high variability in gut microbial populations, though, lending to the validity of this animal model.

In summary, several lines of evidence suggest that cocaine-seeking may be influenced by the gut-brain axis (Kiraly et al., 2016; Meckel and Kiraly, 2019; Chivero et al., 2019; Cryan et al., 2019), and the present work extends this concept by identifying microbial populations that are associated with specific phases of cocaine experience, including candidate biomarkers of cocaine resilience or vulnerability. Given that only a subset of cocaine-experienced individuals develop a substance use disorder (McLellan, 2017), predictive microbiome profiles could suggest new approaches to identifying addiction-prone populations. Moreover, pre-, and probiotic formulas that promote the growth of bacterial species associated with low cocaine vulnerability may ultimately serve as effective adjunct therapies to treat addiction.

## Supplemental Figures

**Supplemental Figure 1:**
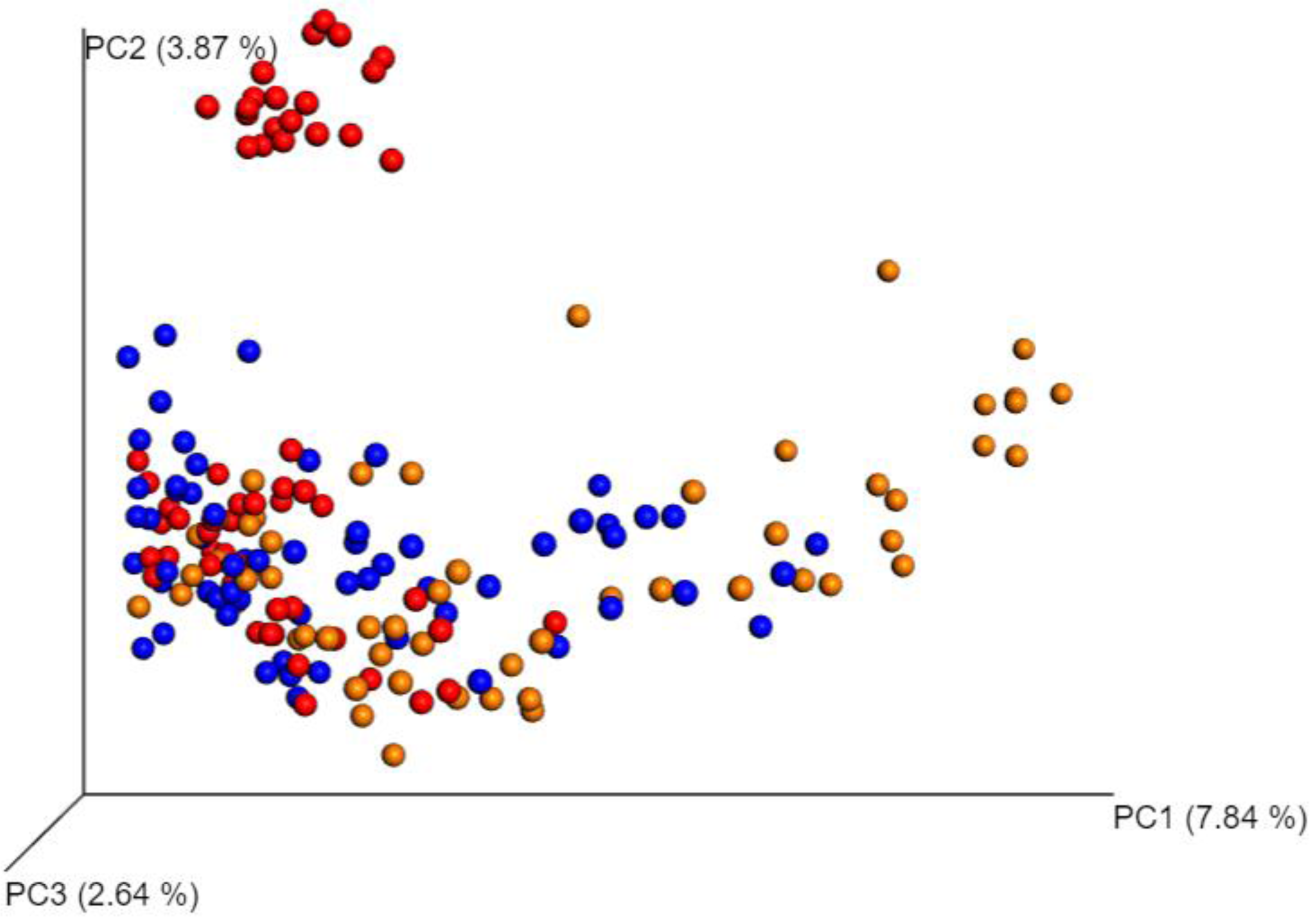
Comparison of cohort influence on microbiota composition. Principal coordinates analysis (PCoA) of the unweighted UniFrac distance matrix for all analyzed fecal samples from all animals, all timepoints, color-coded by cohort (red= cohort 1, blue= cohort 2, gold= cohort 3).

**Supplemental Figure 2:**
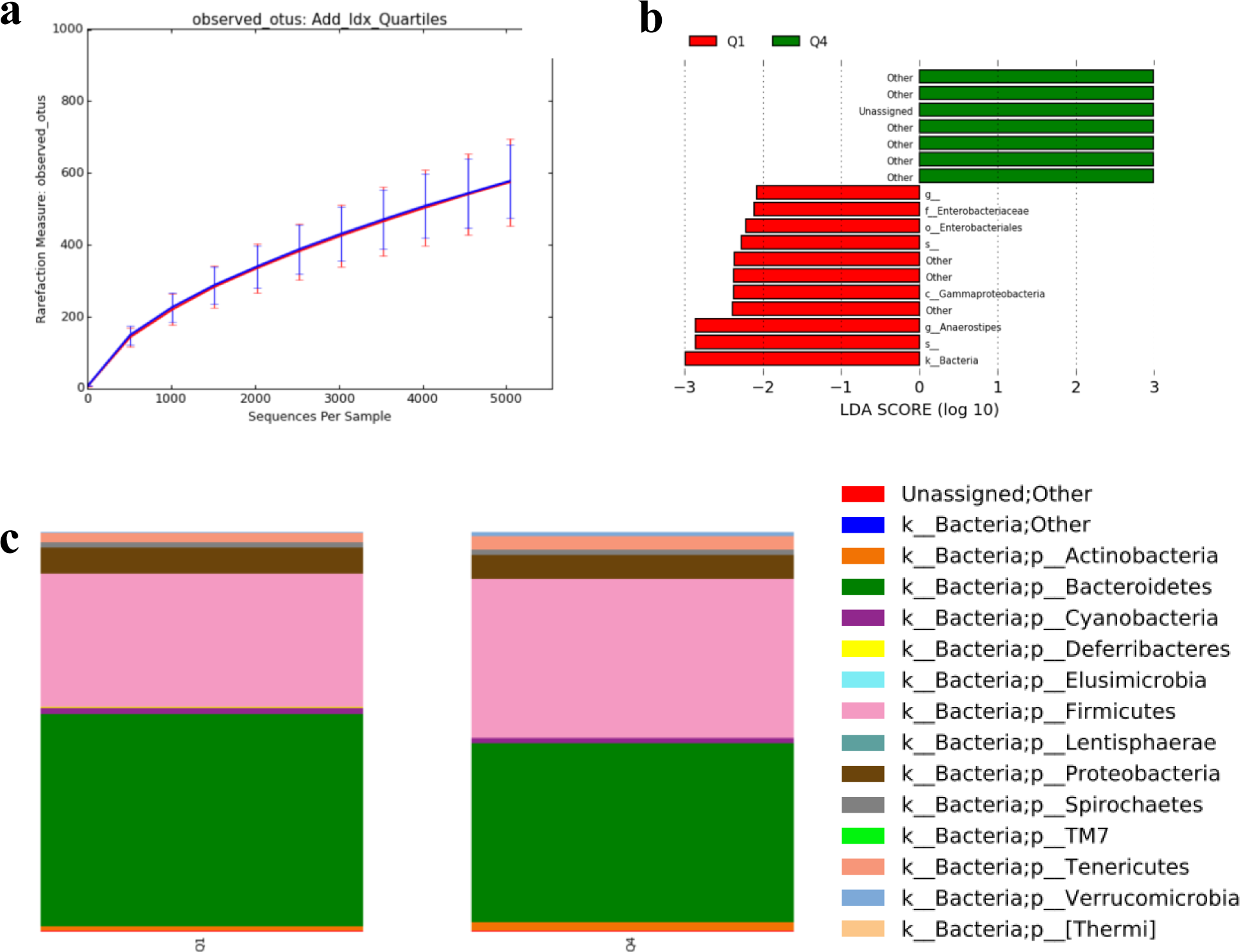
Characterization of microbiota after short access cocaine self-administration in addiction resistant vs addiction prone rats. **a.** α-diversity in fecal samples collected after short-access cocaine self-administration, based on observed OTUs compared between rats categorized as low responders on the overall addiction index (Q1, red) or high responders (Q4, blue). **b.** Linear discriminant analysis effect size was used to investigate whether individual bacterial taxa were more robustly represented in low vs high cocaine responders (Q1, red; Q4, green). **c.** Taxa summarization performed at the phylum level in samples collected after short access self-administration. OTU, operational taxonomic unit; Q, quartile.

**Supplemental Figure 3:**
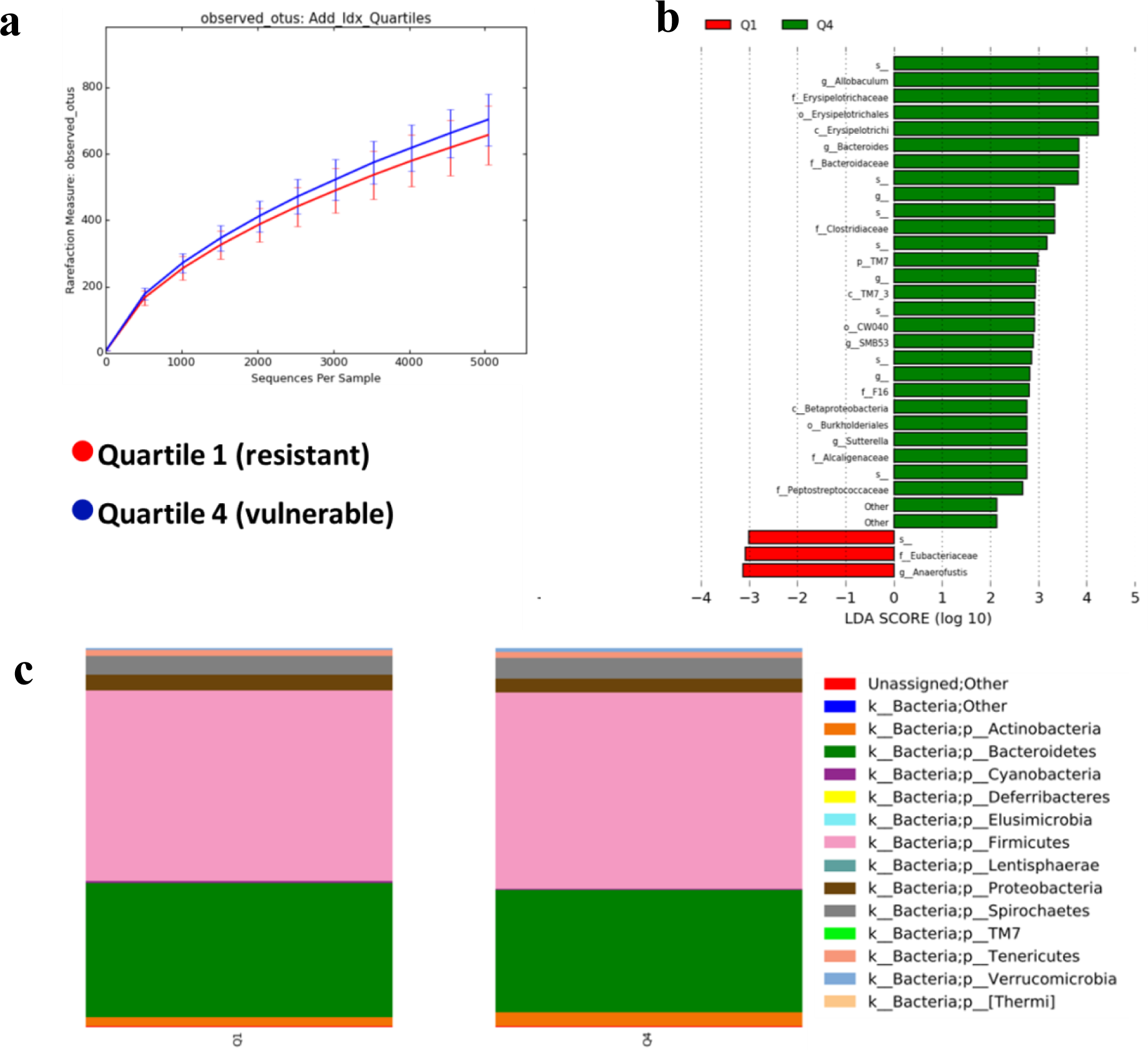
Characterization of microbiota after long access cocaine self-administration in addiction resistant vs addiction prone rats. **a.** α-diversity in fecal samples collected after long access cocaine self-administration, based on observed OTUs compared between rats categorized as low responders on the overall addiction index (Q1, red) or high responders (Q4, blue)**. b.** Linear discriminant analysis effect size was used to investigate whether individual bacterial taxa were more robustly represented in low (red) vs high (green) cocaine responders after prolonged cocaine exposure. **c.** Taxa summarization performed at the phylum level in samples collected after long access self-administration. OTU, operational taxonomic unit; Q, quartile.

**Supplemental Figure 4:**
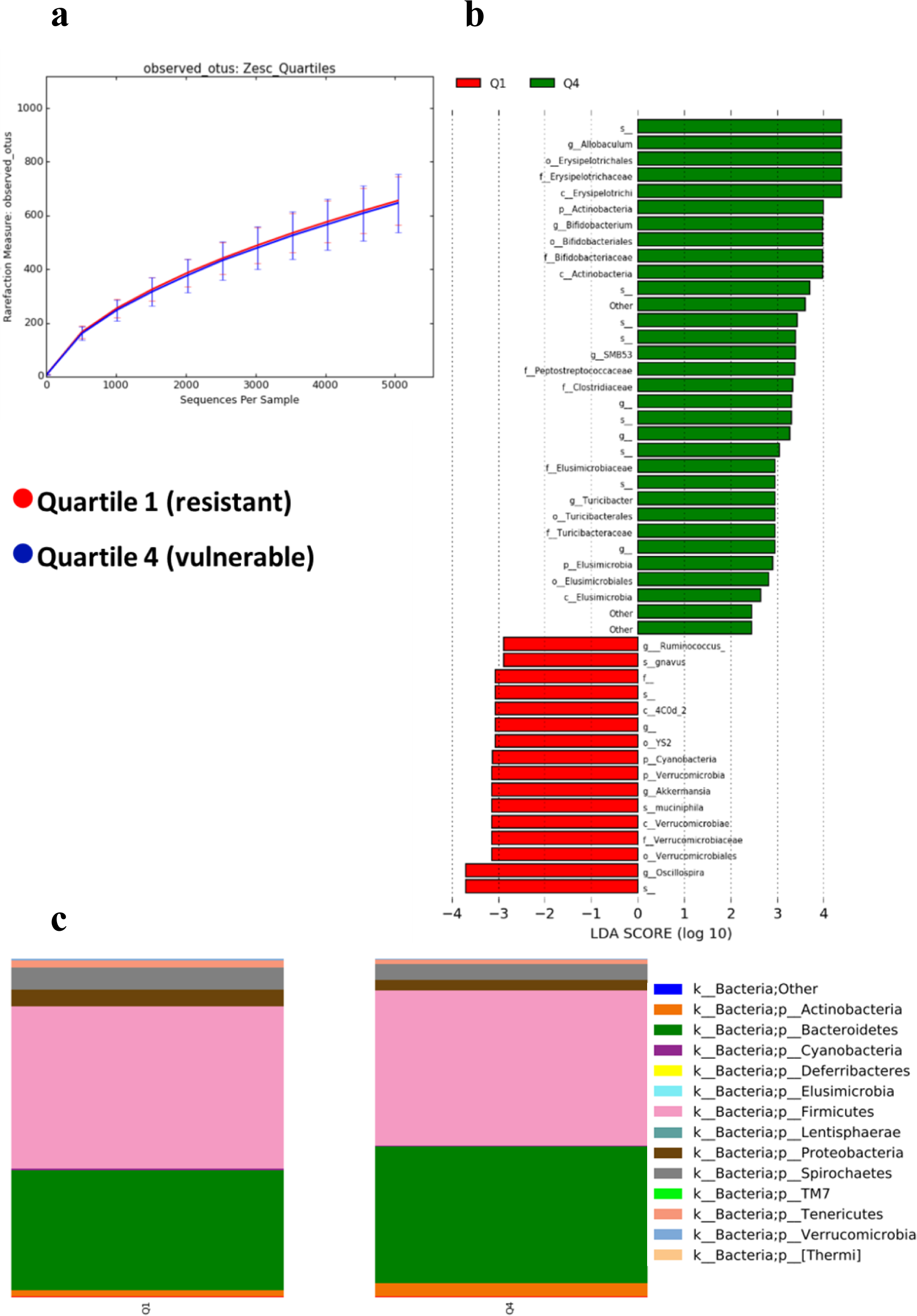
Characterization of microbiota after long access cocaine self-administration in rats with low vs high rates of escalation. **a.** α-diversity in fecal samples collected after long access cocaine self-administration, based on observed OTUs compared between rats categorized as low responders on the escalation index alone (Quartile 1, red) or high responders (Quartile 4, blue)**. b.** Linear discriminant analysis effect size was used to investigate whether individual bacterial taxa were more robustly represented in low vs high cocaine responders after prolonged cocaine exposure using the escalation index to categorize response levels. **c.** Taxa summarization performed at the phylum level in samples collected after long access self-administration using the escalation index to categorize addiction phenotype.

**Supplemental Figure 5:**
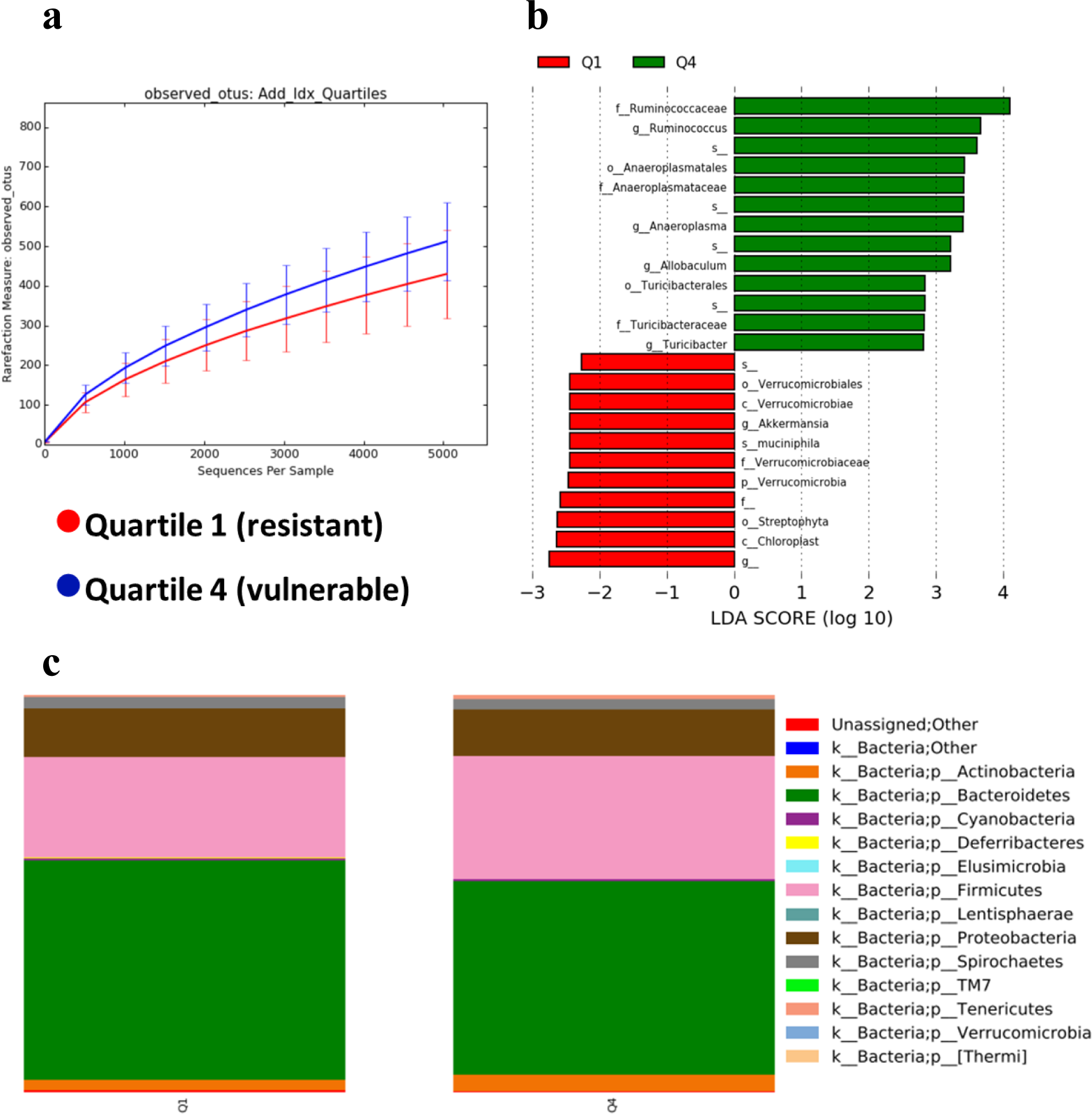
Characterization of microbiota at baseline predicts future cocaine phenotypes. **a.** α-diversity in fecal samples at baseline before cocaine self-administration, based on observed OTUs compared between rats that ultimately categorized as cocaine resistant or vulnerable, based on the overall addiction index. **b.** Linear discriminant analysis effect size was used to investigate bacterial members that might drive differences animals that become low vs high cocaine responders. **c.** Taxa summarization performed at the phylum level. At baseline, there are differences in relative phylum abundance between animals that became low cocaine responders vs high cocaine responders.

**Supplemental Figure 6:**
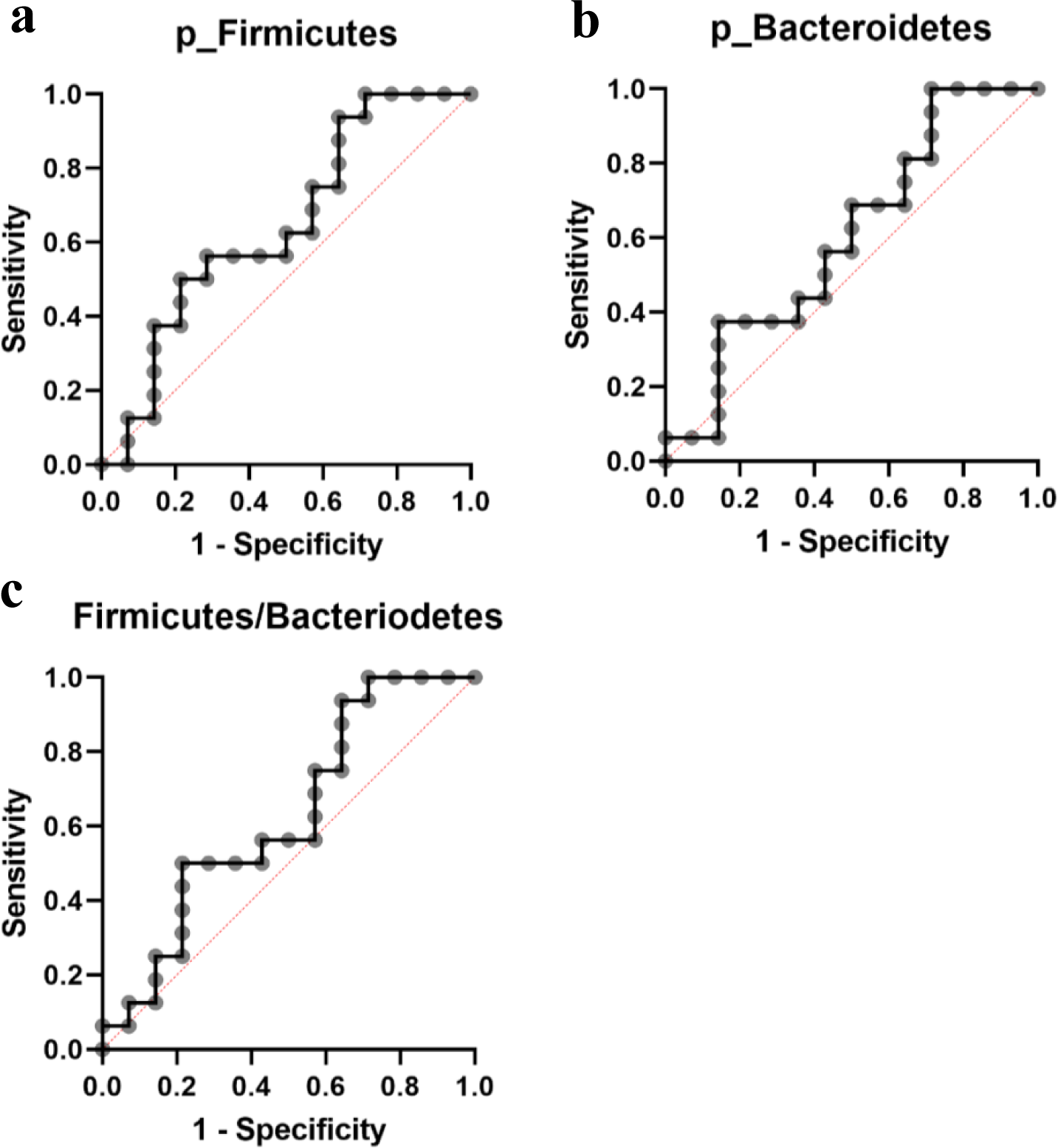
Testing whether the relative abundance of bacterial taxa associated with obesity and gut-related disorders can predict future addiction phenotypes. Receiver operator curves (ROC) showed that baseline relative abundance of the phyla **a.** Firmicutes (AUC=0.6429) and **b.** Bacteroidetes (AUC=0.6027) were not effective at predicting future addiction phenotypes, and **c.** the ratio of these two phyla was not effective either (AUC=0.625).

**Supplemental Figure 7:**
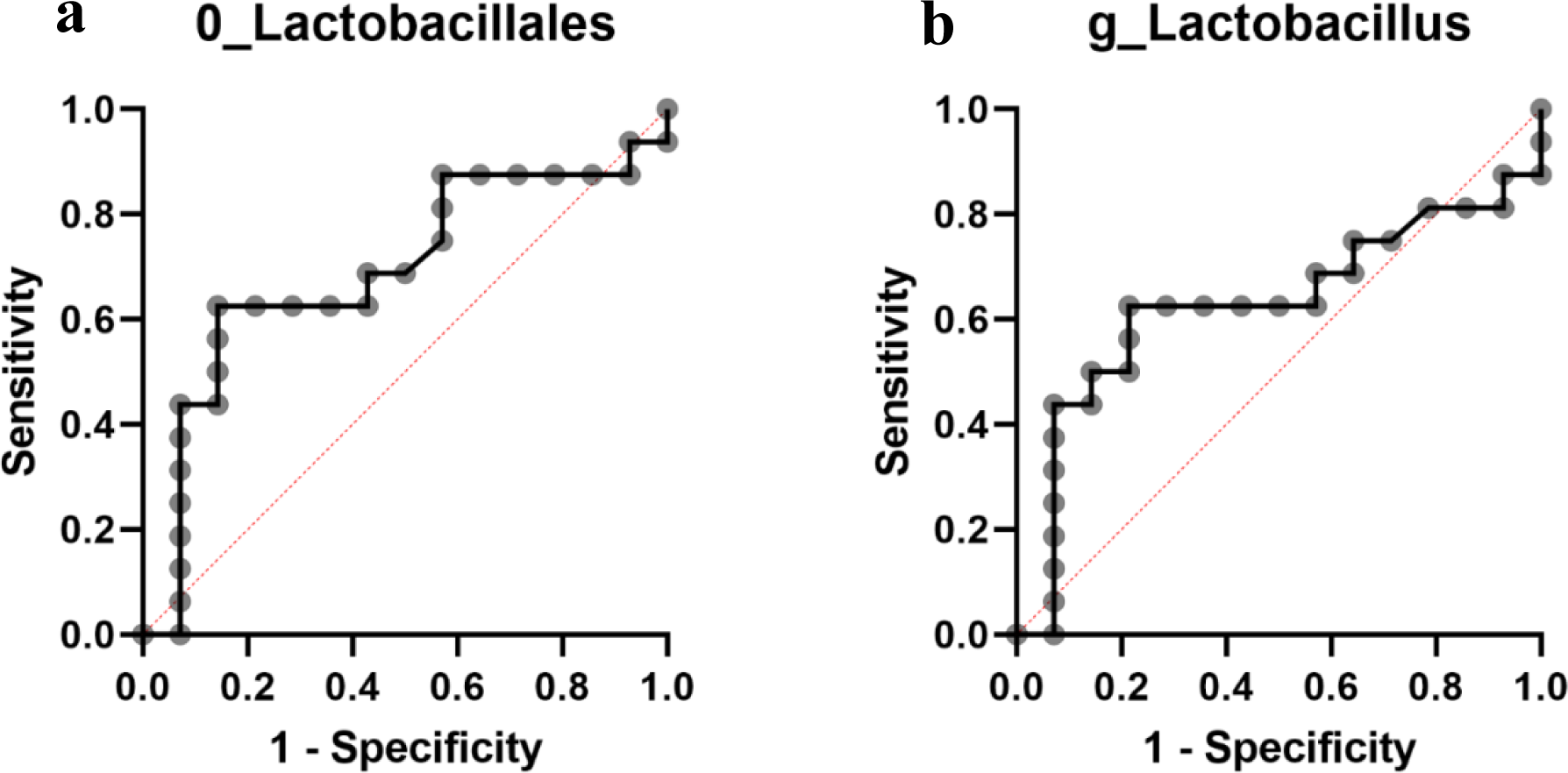
Testing whether the relative abundance of bacterial taxa associated with anxiety- and depression-like behaviors can predict future addiction phenotypes. Receiver operator curves (ROC) showed that baseline relative abundance levels of **a**. the order *Lactobacillales* (AUC=0.6897), and **b**. the genus *Lactobacillus* (AUC=0.6272) were only fair predictors of future addiction phenotypes.

## Acknowledgements

We would like to thank Bonnie Williams, Samantha Spencer, and all other members of the Frantz, Chassaing, and George Labs for project support throughout the study. The Cocaine Biobank is supported by NIH DA043799. KJF’s laboratory is supported by The Center for Behavioral Neuroscience at Georgia State University. GS was a Brains and Behavior Scholar and a graduate assistant with IMSD (R25GM11353206). JSK was supported by IMSD (R25GM11353206). BC’s laboratory is supported by a Starting Grant from the European Research Council (ERC) under the European Union’s Horizon 2020 research and innovation programme (grant agreement No. ERC-2018-StG-804135), a Chaire d’Excellence from IdEx Université de Paris - ANR-18-IDEX-0001, and an Innovator Award from the Kenneth Rainin Foundation. Funders had no role in the design of the study and data collection, analysis and interpretation, nor in manuscript writing.

